# Specification of oxytocinergic and vasopressinergic circuits in the developing mouse brain

**DOI:** 10.1101/2020.08.07.241364

**Authors:** M. Pilar Madrigal Verdú, Sandra Jurado

## Abstract

Oxytocin (OXT) and arginine vasopressin (AVP) support sex-specific and context-appropriate social behaviors. Although alterations of these systems underlie the appearance of neuropsychiatric disorders, their formation and developmental dynamics remain largely unknown. Using novel brain clearing techniques and 3D imaging, we have reconstructed the specification of oxytocinergic and vasopressinergic circuits in the developing mouse brain with unprecedented cellular resolution. A systematic quantification indicates that OXT and AVP neurons in the hypothalamus display distinctive dynamics, but also share common features as a high cellular plasticity from embryonic to early postnatal stages. Our findings reveal new insights into the appearance and consolidation of neuropeptidergic systems in the developing CNS which is a critical step to unveil brain formation and function.

## Introduction

Oxytocin (OXT) and arginine vasopressin (AVP) are evolutionarily conserved neuropeptides implicated in the regulation of complex social behaviors. These neuropeptides are synthesized in the hypothalamus where are processed into a nine amino acid peptide chain only differing in two amino acids^1^. OXT was first identified for its systemic function in parturition and lactation^2-4^. Additionally, cumulative evidence indicated a prominent role of OXT as a pro-social neuromodulator increasing the salience of social stimuli and regulating social behaviors as well as aggression, anxiety, fear, and trust^5-15^. Conversely, AVP has been proposed to antagonize OXT-mediated functions^16^ like increasing social stress^17^ and reducing interpersonal trust in humans^17-19^. However, this view is being revisited after several studies exposed a more complex scenario for the interplay of OXT and AVP in the central nervous system (CNS)^20^. Despite the details of how OXT and AVP interact to modulate neuronal function remain unknown, numerous studies have revealed that alterations of OXT and AVP circuits may underlie mental disorders often characterized by deficits in social interaction such as autism, social anxiety, and schizophrenia^20^.

These data suggest that the balance between OXT- and AVP-mediated signaling is likely to determine the display of appropriate social behaviors, thus understanding their simultaneous developmental dynamics is crucial for having a complete picture of their regulatory roles. Most studies characterizing the expression of OXT and AVP projections have employed histological methods and *in situ* hybridization in fixed sections^21-25^ which provide insightful information, but difficult to extrapolate to circuit formation in the entire brain. Furthermore, most of the previous work has focused on the rat brain^26-32^ despite an increased number of studies employ the mouse as an experimental model, thus increasing the need for accurate connectivity maps for this commonly used species.

To overcome these current challenges, we have implemented the iDISCO^+^ clearing technique to analyze the expression of endogenous neuropeptides. This method in combination with light sheet fluorescent microscopy has allowed us to generate 3D reconstructions of the oxytocinergic and vasopressinergic systems in the entire mouse brain from early development to adulthood. This methodology achieves cell-specific resolution allowing a systematic quantification of oxytocinergic and vasopressinergic cells. Our data revealed that OXT and AVP neurons first appear at the caudal hypothalamic regions followed by more rostral areas. Interestingly, different hypothalamic nuclei show marked differences between OXT and AVP expression during development. The rostral nuclei like the anterodorsal preoptic nucleus (ADPN) and the periventricular nucleus (PeVN) together with neighboring areas like the bed nucleus of strial terminalis (BNST) display a fairly homogeneous expression whereas the caudal nuclei (supraoptic nucleus (SON), paraventricular nucleus (PVN) and the retrochiasmatic nucleus (RCH)) exhibit greater heterogeneity with a large population of neurons coexpressing both OXT and AVP during early postnatal stages. Our analysis indicates that the accessory nucleus (AN) is the most heterogeneous of the hypothalamic nuclei with a rostral section almost exclusively constituted by OXT and OXT+AVP neurons and a caudal region entirely enriched with AVP neurons.

Although the expression of OXT+AVP neurons is quite high at PN7, most of the nuclei experience an increase in the number of OXT neurons in detriment of the mixed OXT+AVP population over postnatal development. This switch in neuropeptide expression is particularly significant in BNST and ADPN, where the presence of AVP neurons decreases drastically in the adult brain. These developmental adaptations are expected to have functional consequences impacting the ratio of AVP/OXT innervation to their projection sites.

In addition to wild type animals, we have characterized OXT expression in a commercial OXT-Cre mouse line^33^. These studies have confirmed our observation of a significant increase of oxytocinergic neurons in most hypothalamic nuclei as development progresses. Strikingly, some OXT positive neurons in the SON and RCH of the commercial OXT-Cre line are not recognized by standard OXT antibodies suggesting the presence of nucleus-specific immature OXT variants.

In summary, our in-depth circuit analysis has revealed that OXT and AVP expression exhibits distinct developmental dynamics in the mouse brain. These dynamic adaptations are likely to modulate the functional properties of different brain regions according to their developmental stage, and thus contributing to the refinement of the neuronal circuits that support context-appropriate social behaviors later in life.

## Results

### Application of iDISCO^+^ for identifying OXT and AVP hypothalamic circuits

Understanding the formation and developmental dynamics of oxytocinergic and vasopressinergic circuits is of great relevance given their regulatory role in social behaviors. The identification of alterations within these systems is expected to reveal critical insights into the underlying causes of several neuropsychiatric disorders^20^. Most of OXT and AVP innervation in the CNS comes from the hypothalamus, a brain area comprised by highly heterogeneous nuclei. Pioneer work using *in situ* hybridization and immunohistochemistry techniques described with great detail the hypothalamic circuit in the developing rat brain^26-32^, however the mouse brain attracted less attention partly because of its smaller size which hinders the identification of the intricate hypothalamic nuclei. Nevertheless, as an increasing number of studies particularly on social behavior utilize mouse models, it is necessary to better understand the development of OXT and AVP circuits in this species.

Recent advances on tissue clearing techniques have enabled the study of neuronal connectivity in the whole brain by means of light sheet microscopy and 3D reconstructions^34-37^. Here, we implemented a protocol for 3D imaging of solvent-cleared brains, adapted from the iDISCO^+^ protocol described in Renier et al. 2016. Incubation time for anti-OXT and anti-AVP antibodies was optimized according to brain size and tissue firmness. Best results were obtained with incubation times of 5 days for E16.5 brains, 10 days for PN0, 2 weeks for PN7 and 3 weeks for adult brains. In our hands, these modifications resulted on fully cleared brains (**Fig. 1a**) with specific staining and low background which allowed to unambiguously distinguish single cells in different hypothalamic nuclei and neighboring brain areas (**Fig. 1b, Supplementary video 1**) including (from rostral to caudal): BNST (bed nucleus of strial terminalis), PeVN (periventricular nucleus), VMPO (ventromedial preoptic nucleus), ADPN (anterodorsal preoptic nucleus), SON (supraoptic nucleus), SCH (suprachiasmatic nucleus), PVN (paraventricular nucleus), RCH (retrochiasmatic nucleus) and AN (accessory nucleus, constituted by scattered cells within the hypothalamic area not included in any of the other nuclei^38^). This method in combination with 3D image processing enabled to isolate single hypothalamic nuclei (surface Imaris tool; **Fig. 1c**), and to analyze different neuronal populations within each one (spots Imaris tool; **Fig. 1d**). This technique also allowed us to identify a small population of OXT and AVP positive cells in other brain areas like the medial amygdala (MeA), where these cells were mainly found in the anteriodorsal and anteroventral part (**Table 1**). These results indicate that iDISCO^+^ provides an excellent method to preserve the identity of small protein-like molecules like OXT and AVP, and permits identifying intricate nuclei such as hypothalamic structures.

**Table 1.** Total number of OXT, AVP and OXT+AVP neurons in the developing mouse brain. Quantification of total number of OXT, AVP and OXT+AVP neurons at different developmental stages (E16.5, PN0, PN7 and adult brain). Abbreviations: PeVN, periventricular nucleus; ADPN, anterodorsal preoptic nucleus; BNST, bed nucleus of strial terminalis; MeA, medial amygdala; VMPO, ventromedial preoptic nucleus; SON, supraoptic nucleus; PVN, paraventricular nucleus; RCH, retrochiasmatic nucleus; AN, accessory nucleus; SCH, suprachiasmatic nucleus. Results are expressed as mean ± S.E.M. (n = 4).

| OXT | E16.5 | PN0 | PN7 | Adult |
| --- | --- | --- | --- | --- |
| BNST | 0.00 | 2.00 ± 2.00 | 16.00 ± 9.94 | 100.00 ± 36.46 |
| PVN | 24.50 ± 14.22 | 123.75 ± 33.62 | 131.25 ± 70.70 | 461.25 ± 20.60 |
| SON | 116.75 ± 12.60 | 33.00 ± 13.37 | 83.50 ± 50.49 | 216.00 ± 44.67 |
| VMPO | 0.00 | 6.25 ± 2.46 | 3.25 ± 3.25 | 8.75 ± 4.84 |
| RCH | 99.25 ± 11.92 | 63.25 ± 16.85 | 70.75 ± 8.62 | 101.75 ± 28.05 |
| ADPN | 0.00 | 0.00 | 17.75 ± 14.79 | 75.00 ± 28.67 |
| PeVN | 0.00 | 30.25 ± 18.22 | 50.50 ± 31.62 | 123.50 ± 45.17 |
| AN | 0.00 | 37.5 ± 15.76 | 11.50 ± 5.45 | 164.75 ± 18.28 |
| SCH | 0.00 | 3.67 ± 1.61 | 0.67 ± 0.58 | 4.00 ± 1.78 |
| MeA | 0.00 | 0.25 ± 0.25 | 0.25 ± 0.25 | 0.50 ± 0.50 |

| AVP | E16.5 | PN0 | PN7 | Adult |
| --- | --- | --- | --- | --- |
| BNST | 0.00 | 2.75 ± 1.80 | 9.50 ± 3.80 | 1.5 ± 0.65 |
| PVN | 61.5 ± 8.87 | 138.25 ± 33.69 | 88.00 ± 59.30 | 206.50 ± 74.77 |
| SON | 129 ± 6.36 | 122.25 ± 12.64 | 207.25 ± 17.67 | 169.50 ± 45.75 |
| VMPO | 0.00 | 4.00 ± 0.58 | 2.00 ± 1.22 | 2.25 ± 0.48 |
| RCH | 107.50 ± 20.23 | 73.00 ± 14.70 | 128.00 ± 40.19 | 69.75 ± 8.54 |
| ADPN | 0.00 | 3.25 ± 2.93 | 3.25 ± 2.29 | 2.00 ± 0.71 |
| PeVN | 0.00 | 11.50 ± 4.33 | 56.5 ± 31.47 | 14.5 ± 3.28 |
| AN | 0.00 | 85.00 ± 10.87 | 92.00 ± 16.91 | 95.00 ± 12.51 |
| SCH | 0.00 | 368.67 ± 24.50 | 322.67 ± 27.57 | 175.25 ± 27.68 |
| MeA | 0.00 | 0.00 | 2.75 ± 1.38 | 0.00 |

**Table 1. Total number of OXT, AVP and OXT+AVP neurons in the hypothalamus nuclei during development.**
| OXT-AVP | E16.5 | PN0 | PN7 | Adult |
| --- | --- | --- | --- | --- |
| BNST | 0.00 | 53.75 ± 8.09 | 157.50 ± 11.91 | 53.50 ± 49.58 |
| PVN | 130.75 ± 16.27 | 68.75 ± 31.07 | 388.00 ± 72.50 | 93.50 ± 24.13 |
| SON | 13.50 ± 2.72 | 51.50 ± 18.31 | 168.75 ± 40.24 | 78.75 ± 29.56 |
| VMPO | 0.00 | 8.00 ± 2.80 | 14.50 ± 3.07 | 21.00 ± 12.92 |
| RCH | 11.75 ± 1.25 | 63.75 ± 23.67 | 86.00 ± 32.54 | 50.00 ± 4.42 |
| ADPN | 0.00 | 23.50 ± 6.96 | 58.75 ± 23.57 | 17.00 ± 14.02 |
| PeVN | 0.00 | 10.25 ± 4.25 | 91.00 ± 14.87 | 57.00 ± 37.41 |
| AN | 0.00 | 24.50 ± 14.15 | 148.25 ± 11.52 | 15.50 ± 4.17 |
| SCH | 0.00 | 0.67 ± 0.58 | 2.33 ± 0.29 | 3.00 ± 1.47 |
| MeA | 0.00 | 0.50 ± 0.50 | 7.50 ± 1.19 | 3.00 ± 1.78 |

**Fig. 1.**
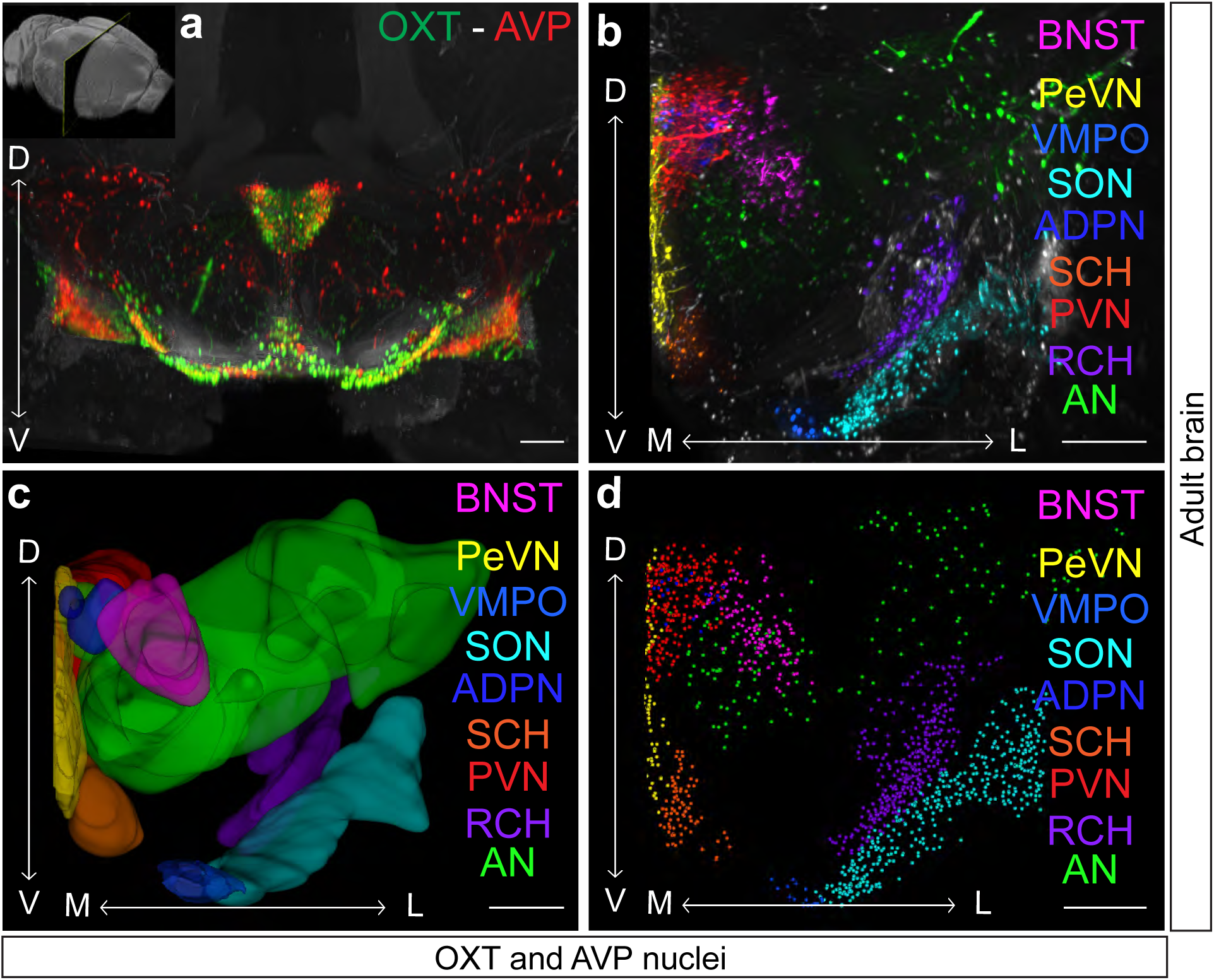
iDISCO^+^, a useful tool for identifying specific hypothalamic and other brain nuclei. Representative images of an adult mouse brain processed with the iDISCO^+^ protocol and stained against OXT (green) and AVP (red) (**a**) in which specific neuronal populations can be easily identified. High magnification shows the hypothalamus of one hemisphere displaying oxytocinergic and vasopressinergic neurons pseudocolored in each hypothalamic nucleus and surrounding areas like the BNST (**b**). OXT and AVP expression in each nucleus are represented using the Imaris surface tool (**c**). OXT and AVP neurons in each nucleus are identified and outlined to individual spots for performing quantitative analysis (**d**). Arrows: D, dorsal; V, ventral; M, medial; L, lateral. Abbreviations: PeVN, periventricular nucleus; ADPN, anterodorsal preoptic nucleus; BNST, bed nucleus of strial terminalis; VMPO, ventromedial preoptic nucleus; SON, supraoptic nucleus; PVN, paraventricular nucleus; RCH, retrochiasmatic nucleus; AN, accessory nucleus; SCH, suprachiasmatic nucleus. Scale bar: 300 µm.

### Distinct hypothalamic nuclei exhibit different temporal windows of OXT and AVP expression

The precise anatomical resolution achieved by iDISCO^+^ allowed to isolate individual hypothalamic nuclei for quantification of both OXT and AVP neurons at different developmental stages (see **Supplementary videos 2-5** for 3D whole-brain reconstructions). Our analysis revealed significant differences in the expression dynamics of OXT and AVP in the developing hypothalamus, from embryo to adulthood. Whereas OXT and AVP positive neurons can be identified at caudal nuclei like the PVN, SON and RCH as early as E16.5, rostral hypothalamic nuclei remained unlabeled (**Fig. 2a-d, Supplementary Fig. 1, Tables 1** and **2**). However, at PN0 all the hypothalamic nuclei showed OXT and AVP neurons, with the exception of the SCH which expresses AVP neurons almost exclusively^39^ (**Fig. 2e-h, Supplementary Fig. 1**). OXT and AVP expression patterns were quite consistent in all hypothalamic nuclei from early postnatal to adulthood (**Fig. 2i-p**). These data indicate that OXT and AVP neurons first appear in caudal areas in agreement with the role of these nuclei as main sources of OXT and AVP modulation (**Fig. 2, Supplementary Fig. 1**)^40^. As a general rule, the number of OXT and AVP neurons rises with development in all hypothalamic areas (**Table 1**). This proliferation is likely to occur in parallel with brain growth and its increasing demands for neuromodulatory innervation.

**Fig. 2.**
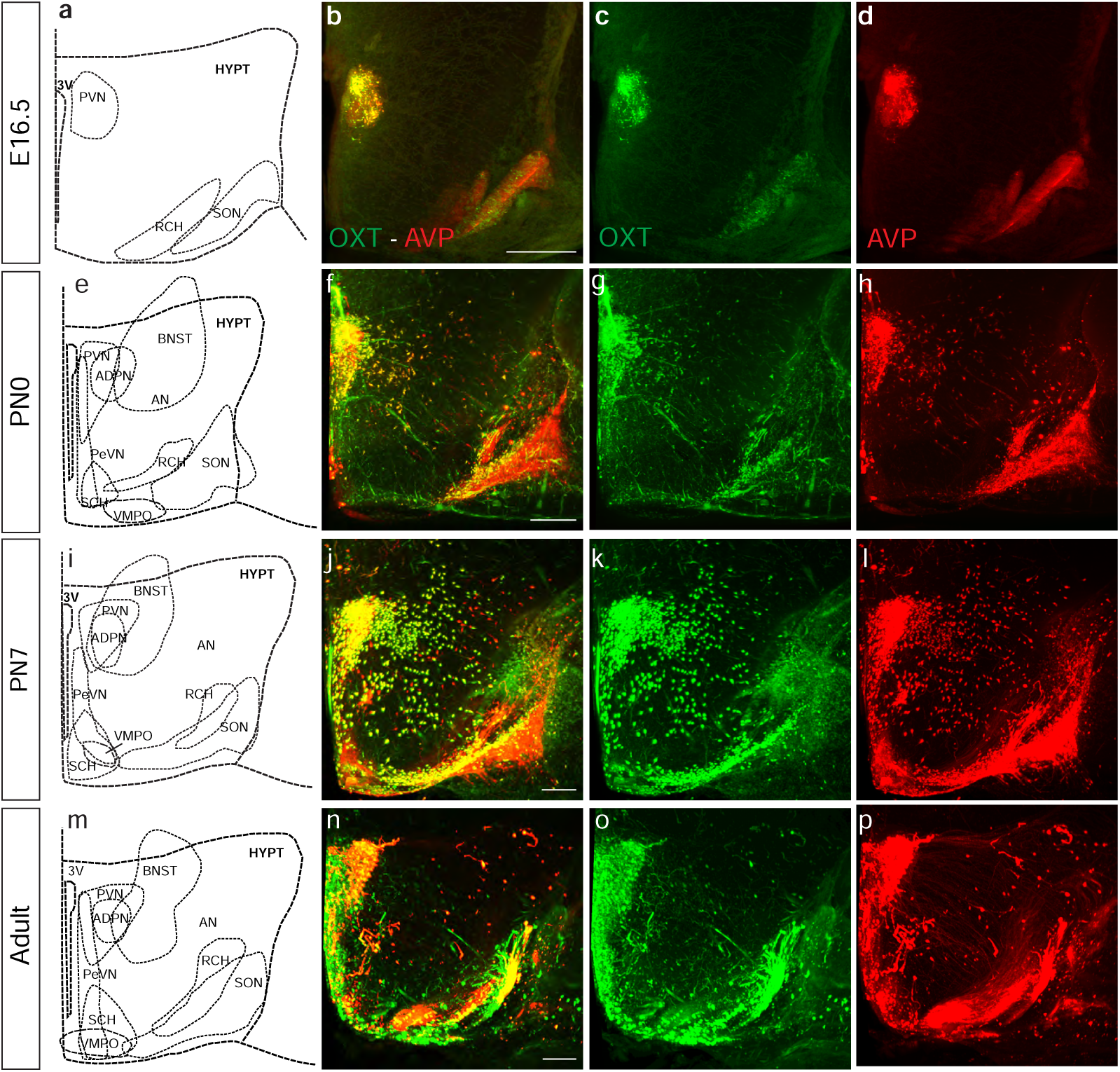
OXT and AVP expression patterns in the developing mouse hypothalamus. Snap shot of a whole brain stained with an anti-OXT (green) and anti-AVP (red) antibody. Brain coronal sections at different developmental stages: E16.5 (**a-d**), PN0 (**e-h**), PN7 (**i-l**) and young adult (**m-p**). Abbreviations: 3v, third ventricle; PeVN, periventricular nucleus; ADPN, anterodorsal preoptic nucleus; BNST, bed nucleus of strial terminalis; VMPO, ventromedial preoptic nucleus; SON, supraoptic nucleus; PVN, paraventricular nucleus; RCH, retrochiasmatic nucleus; AN, accessory nucleus; SCH, suprachiasmatic nucleus; MeA, medial amygdale nucleus. Scale bar: 200 µm.

Temporal and spatial heterogeneity of OXT and AVP expression in different hypothalamic nuclei was highlighted by flatmap representations of 3D brain reconstructions (**Fig. 3**). Hypothalamic flatmaps convey the intensity of OXT and AVP positive cells providing a quantitative representation of cell density in each nucleus. 3D-based hypothalamic flatmaps showed clear differences in the expression pattern of OXT and AVP during development (see density quantifications in **Supplementary Fig. 2**). Remarkably, whereas the density of OXT positive neurons seem to steadily increase (**Fig. 3e-h**), AVP neurons show the opposite behavior (**Fig. 3i-l**). Interestingly, neurons co-expressing OXT and AVP show an increase at early postnatal stage (PN0) (**Fig. 3m-p**) coinciding with a critical period for the maturation of social behaviors.

**Fig. 3.**
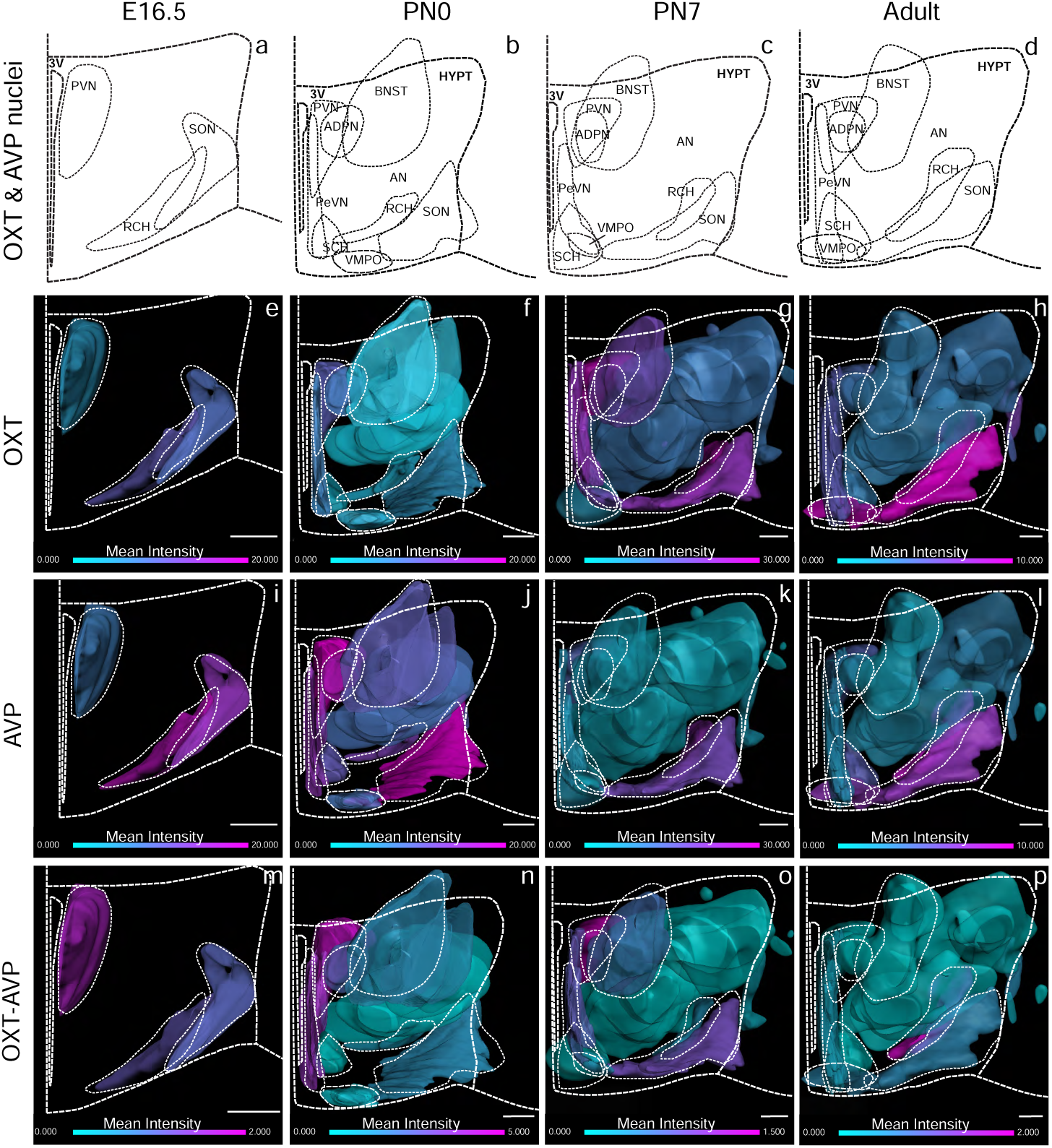
OXT, AVP and OXT+AVP cell density in the developing mouse hypothalamus. 3D hypothalamic flatmap representations of OTX (**e-h**), AVP (**i-l**) and OXT+AVP cells (**m-p**) in each nucleus at different developmental stages: E16.5 (**a**), PN0 (**b**), PN7 (**c**) and adult (**d**). Flatmaps show the expression intensity of each cell population at each developmental stage. Please note the difference intensity scale according to the developmental stage. The minimum intensity is represented in turquoise and the maximum in pink. Abbreviations: 3v, third ventricle; PeVN, periventricular nucleus; ADPN, anterodorsal preoptic nucleus; BNST, bed nucleus of strial terminalis; VMPO, ventromedial preoptic nucleus; SON, supraoptic nucleus; PVN, paraventricular nucleus; RCH, retrochiasmatic nucleus; AN, accessory nucleus; SCH, suprachiasmatic nucleus; MeA, medial amygdale nucleus. Scale bar: 200 µm.

### Neurons co-expressing AVP and OXT are a common feature of the developing hypothalamus

Cell quantification confirmed that AVP and OXT systems in different hypothalamic nuclei differ in their temporal dynamics (**Fig. 4**). The percentage of AVP neurons is higher earlier in life and progressively declines during adulthood in most of the nuclei (**Figs. 4** and **5, Table 2**, percentage of AVP neurons normalized to total number of cells: E16.5: 42.62 ± 2.78; PN0: 52.25 ± 6.04; PN7: 27.97 ± 3.11; Adult: 27.43 ± 3.65; n=4 per each developmental stage, mean ± S.E.M). This reduction is also observed in AVP-dominant nuclei like the SCH^39^ where the percentage of AVP-expressing cells decreases from early postnatal stages (PN0) to adulthood (**Fig. 4i’’**; and **5**). Most nuclei show a similar reduction in the percentage of AVP cells with maturation, with the exception of the PVN that exhibits a non-significant increase vasopressinergic neurons in the adult brain (**Fig. 4e’’, Table 2**).

**Table 2.** Percentage of OXT, AVP and OXT+AVP neurons in the developing mouse brain. Quantification of the percentage of each neuron population at different developmental stages (E16.5, PN0, PN7 and adult brain). Abbreviations: PeVN, periventricular nucleus; ADPN, anterodorsal preoptic nucleus; BNST, bed nucleus of strial terminalis; VMPO, ventromedial preoptic nucleus; SON, supraoptic nucleus; PVN, paraventricular nucleus; RCH, retrochiasmatic nucleus; AN, accessory nucleus; SCH, suprachiasmatic nucleus. Results are expressed as mean ± S.E.M. (n = 4).

| **OXT** | **E16.5** | **PN0** | **PN7** | **Adult** |
| --- | --- | --- | --- | --- |
| **BNST** | 0.00 | 4.35 ± 4.35 | 7.418 ± 4.41 | 72.63 ± 24.28 |
| **PVN** | 10.65 ± 5.94 | 35.08 ± 7.33 | 19.54 ± 9.38 | 61.76 ± 4.29 |
| **SON** | 44.81 ± 3.16 | 15.79 ± 6.52 | 17.16 ± 9.96 | 45.06 ± 4.46 |
| **VMPO** | 0.00 | 34.29 ± 12.42 | 8.13 ± 8.13 | 41.55 ± 22.95 |
| **RCH** | 46.30 ± 5.70 | 30.91 ± 8.00 | 29.87 ± 7.97 | 43.84 ± 7.87 |
| **ADPN** | 0.00 | 0.00 | 18.40 ± 7.25 | 71.26 ± 23.78 |
| **PeVN** | 0.00 | 49.41 ± 15.75 | 17.90 ± 10.50 | 61.42 ± 20.64 |
| **AN** | 0.00 | 23.49 ± 9.16 | 4.35 ± 2.02 | 59.79 ± 1.68 |
| **SCH** | 0.00 | 0.69 ± 0.40 | 0.16 ± 0.16 | 2.22 ± 0.86 |
| **MeA** | 0.00 | 25.00 ± 25.00 | 0.00 | 25.00 ± 25.00 |

| **AVP** | **E16.5** | **PN0** | **PN7** | **Adult** |
| --- | --- | --- | --- | --- |
| **BNST** | 0.00 | 5.49 ± 4.02 | 4.90 ± 1.77 | 0.82 ± 0.33 |
| **PVN** | 28.23 ± 3.72 | 39.63 ± 7.15 | 12.30 ± 7.02 | 25.26 ± 6.12 |
| **SON** | 50.03 ± 2.70 | 58.93 ± 3.12 | 45.38 ± 2.49 | 34.45 ± 6.12 |
| **VMPO** | 0.00 | 22.80 ± 4.63 | 10.63 ± 7.10 | 8.10 ± 2.18 |
| **RCH** | 48.15 ± 5.82 | 38.04 ± 8.18 | 42.49 ± 4.61 | 32.74 ± 5.28 |
| **ADPN** | 0.00 | 11.46 ± 7.86 | 3.69 ± 1.90 | 2.24 ± 1.02 |
| **PeVN** | 0.00 | 21.90 ± 5.20 | 22.78 ± 8.09 | 7.55 ± 1.88 |
| **AN** | 0.00 | 57.53 ± 5.01 | 36.18 ± 6.04 | 34.62 ± 2.60 |
| **SCH** | 0.00 | 98.93 ± 0.48 | 99.08 ± 0.14 | 96.05 ± 1.22 |
| **MeA** | 0.00 | 0.00 | 22.69 ± 8.60 | 0.00 |

**Table 2. Percentage of OXT, AVP and OXT+AVP neurons in the hypothalamus nuclei during development.**
| **OXT+AVP** | **E16.5** | **PN0** | **PN7** | **Adult** |
| --- | --- | --- | --- | --- |
| **BNST** | 0.00 | 90.16 ± 8.34 | 87.68 ± 5.95 | 26.55 ± 24.07 |
| **PVN** | 61.12 ± 8.71 | 25.29 ± 14.44 | 68.16 ± 15.94 | 12.98 ± 4.06 |
| **SON** | 5.16 ± 0.93 | 25.28 ± 9.57 | 38.19 ± 9.90 | 20.48 ± 9.86 |
| **VMPO** | 0.00 | 42.92 ± 14.30 | 81.25 ± 11.25 | 50.35 ± 25.07 |
| **RCH** | 5.56 ± 0.75 | 31.06 ± 10.80 | 27.64 ± 3.78 | 23.43 ± 3.36 |
| **ADPN** | 0.00 | 88.54 ± 7.86 | 77.91 ± 8.45 | 26.50 ± 22.93 |
| **PeVN** | 0.00 | 28.69 ± 11.73 | 59.33 ± 18.03 | 31.03 ± 21.02 |
| **AN** | 0.00 | 18.98 ± 11.00 | 59.47 ± 6.30 | 5.59 ± 1.34 |
| **SCH** | 0.00 | 0.12 ± 0.12 | 0.54 ± 0.18 | 1.73 ± 0.79 |
| **MeA** | 0.00 | 25.00 ± 25.00 | 52.31 ± 17.91 | 50.00 ± 28.87 |

**Fig. 4.**
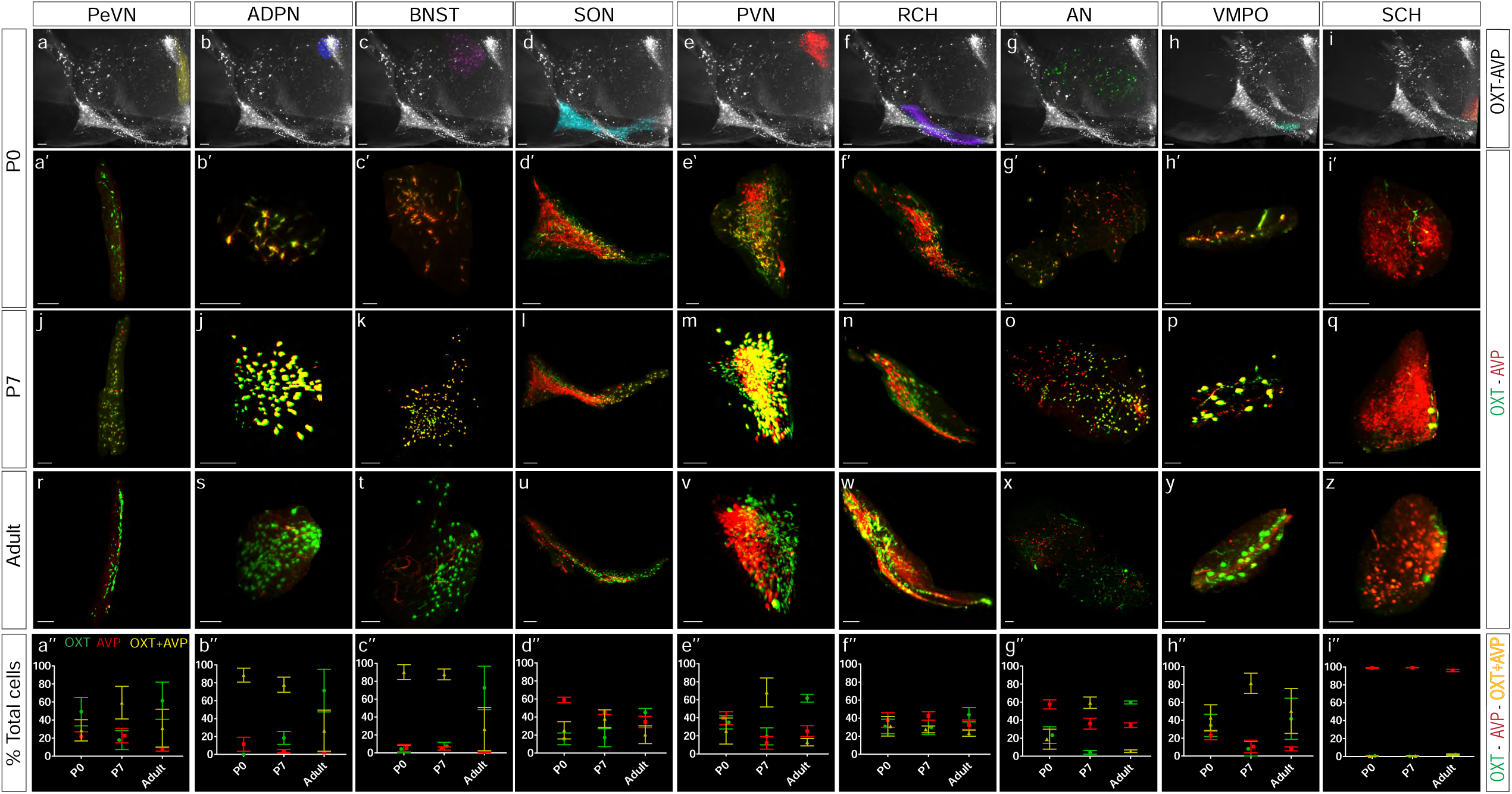
Different nuclei exhibit distinct OXT and AVP expression dynamics during development. Whole-mount immunolabeling against OXT and AVP at PN0 mouse hypothalamus pseudocolored in each nucleus (**a-i**). Nucleus segmentation shows OXT (green) and AVP (red) at different stages: PN0 (**a’-i’**), PN7 (**j-q**), and young adult (**r-z**). Percentage of each neuronal population, OXT, AVP and OXT+AVP neurons, in each nucleus during development (**a’’-i’’;** *n* = 4; data are represented as mean ± S.E.M). Abbreviations: PeVN, periventricular nucleus; ADPN, anterodorsal preoptic nucleus; BNST, bed nucleus of strial terminalis; VMPO, ventromedial preoptic nucleus; SON, supraoptic nucleus; PVN, paraventricular nucleus; RCH, retrochiasmatic nucleus; AN, accessory nucleus; SCH, suprachiasmatic nucleus. Scale bar: 100 µm.

3D imaging quantification revealed that at early postnatal stages most hypothalamic nuclei exhibit a high percentage of neurons co-expressing OXT and AVP (**Figs. 4a’’-i’** and **5, Tables 1** and **2**; percentage of cells from the whole hypothalamus combining all nuclei E16.5: OXT = 34.66 % ± 3.21; AVP = 42.62 % ± 2.78; OXT+AVP = 22.72 % ± 2.75; PN0: OXT = 20.93 % ± 4.91; AVP = 52.25 % ± 6.04; OXT+AVP = 26.83 % ± 10.74; PN7: OXT = 12.91 % ± 4.41; AVP = 27.97 % ± 3.11; OXT+AVP = 39.39 % ± 6.97; Adult: OXT = 52.00 % ± 5.10; AVP = 27.43 % ± 3.65; OXT+AVP = 17.25 % ± 5.91; n=4 per each stage). This phenomenon is quite prominent in some nuclei such as ADPN and neighboring areas like the BNST, where the percentage of neurons co-expressing OXT and AVP decreases in the adult brain, although in a rather heterogeneous fashion (**Figs. 4b’’, 4c’’, 5** and **Table 2**, percentage of AVP+OXT neurons: PN0: ADPN = 88.54 % ± 7.86; BNST = 90.16 % ± 8.34; PN7: ADPN = 77.91 % ± 8.45; BNST = 87.68 % ± 5.95; Adult: ADPN = 26.50 % ± 22.93; BNST = 26.55 % ± 24.07; n=4 per each stage). In the rest of the nuclei, co-labeling of OXT and AVP reaches a peak at PN7 that steadily decreases over time (**Fig. 4a’’-h’’**), with the exception of SCH which is primarily constituted by AVP-expressing neurons (**Figs. 4i’’** and **5, Tables 1** and **2**; SCH neurons: PN0: OXT: 0.69% ± 0.40; AVP = 98.93 % ± 0.48; OXT+AVP = 0.12 % ± 0.12; PN7: OXT = 0.16 % ± 0.16; AVP = 99.08 % ± 0.14; OXT+AVP = 0.54 % ± 0.18; Adult: OXT = 2.22 % ± 0.86; AVP = 96.05% ± 1.22; OXT+AVP = 1.73 % ± 0.79; n=4 per each stage).

**Fig. 5.**
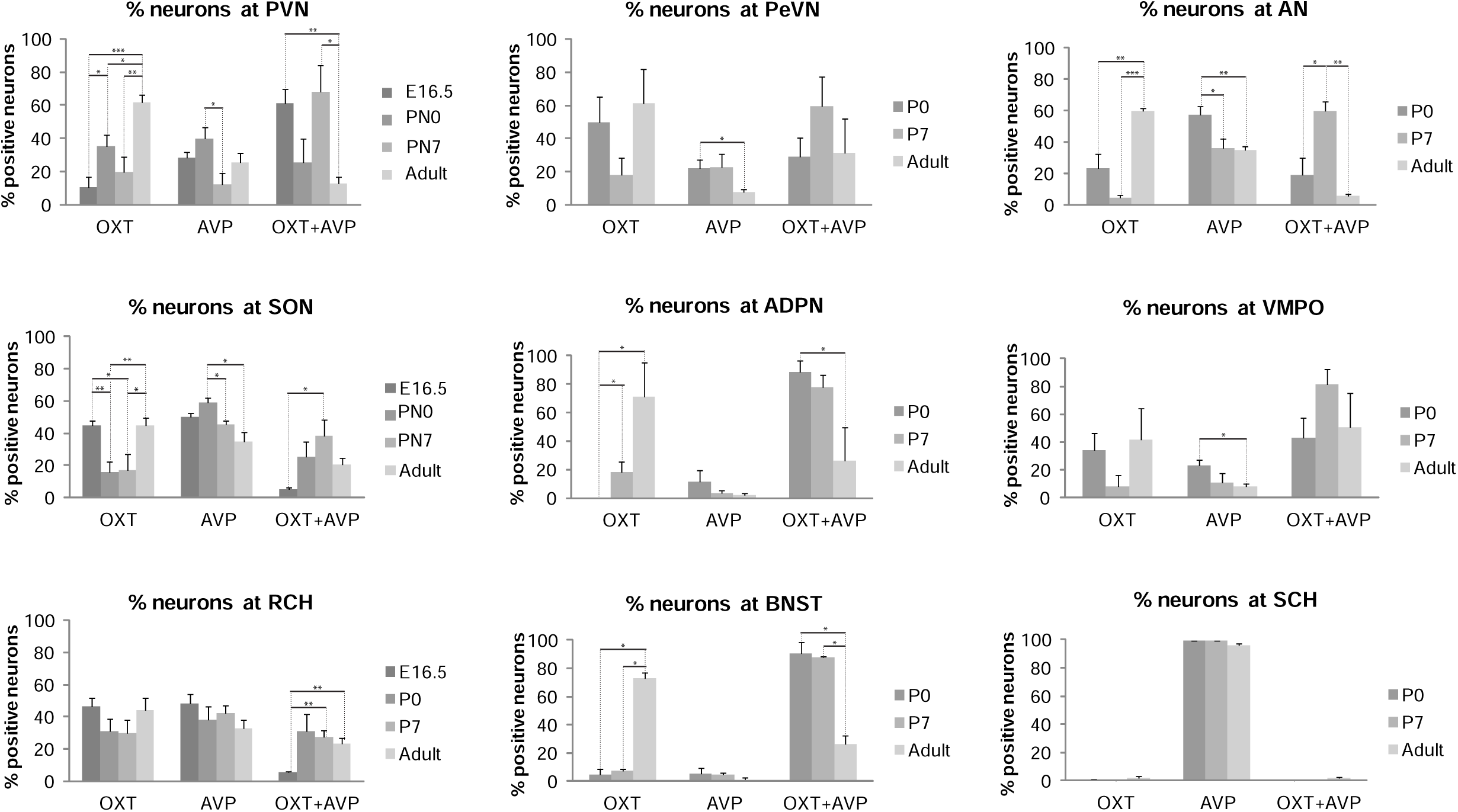
Quantification of OXT and AVP neurons during development. Percentage of OXT, AVP and OXT+AVP neurons is represented at E16.5, PN0, PN7 and adult brain. OXT and AVP co-labeling peaks at PN7 and then decreases in favor of OXT expression in most nuclei with the exception of SCH which is primarily constituted by AVP-expressing neurons. Data are represented as mean ± S.E.M (n = 4). *T-Student* test, *p* – value % OXT: PVN_E16.5-P0_ = 0.041; PVN_E16.5-Adult_ = 0,00043; PVNP0-Adult = 0,02003; pVNP7-Adult = 0,00640; SON_E16.5-P0_ = 0,00706; SON_E16.5-P7_ = 0,03818; SON_P0-Adult_ = 0,00999; SON_P7-Adult_ = 0,04302; ADPN_P0-P7_ = 0,04423; ADPN_P0-Adult_ = 0,02410; AN _P0-Adult_ = 0,00800; AN _P7-Adult_ = 0,000001; *p* – value % AVP: PVN_P0-P7_ = 0,03430; SON_P0-P7_ = 0,01465; SON_P0-Adult_ = 0,01190; VMPO_P0-Adult_ = 0,02835; AN_P0-P7_ = 0,03459; AN_P0-Adult_ = 0,00668; *p* – value % OXT+AVP: BNST_P0-Adult_ = 0,04666; BNST_P7-Adult_ = 0,04869; PVN_E16.5-Adult_ = 0,00243; PVN_P7-Adult_ = 0,01533; SON_E16.5-P7_ = 0,01596; RCH_E16.5-P7_ = 0,00122; RCH_E16.5-Adult_ = 0,00205; ADPN_P0-Adult_ = 0,04295; AN_P0-P7_ = 0,01873; AN_P7-Adult_ = 0,00016. Abbreviations: PeVN, periventricular nucleus; ADPN, anterodorsal preoptic nucleus; BNST, bed nucleus of strial terminalis; VMPO, ventromedial preoptic nucleus; SON, supraoptic nucleus; PVN, paraventricular nucleus; RCH, retrochiasmatic nucleus; AN, accessory nucleus; SCH, suprachiasmatic nucleus.

In contrast, OXT neurons exhibit the opposite trend with their lowest expression at PN7 from when OXT significantly rises to reach maximum levels in the adult brain (**Fig. 4**). The transition from OXT+AVP to OXT expression is more pronounced in the BNST and ADPN where the percentage of AVP neurons is drastically reduced in the adult brain, although with some variability^24^ (**Fig. 4b’’** and **4c’’**). These results indicate that neurons co-expressing OXT and AVP are a common feature of the developing mouse hypothalamus. The functional role of this mixed neuronal population remains unexplored, however it is tempting to speculate that OXT and AVP may share common signaling pathways during early development as immature forms of the OXT and AVP receptors may be activated by both neuropeptides during embryonic and early postnatal stages^41,42^. Furthermore, our results highlight that OXT and AVP neurons follow independent specification patterns in different hypothalamic nuclei. Together, these data show that OXT and AVP systems in the mouse brain are highly dynamic during early development and continuing their remodeling until well into the postnatal period.

### Cell diversity of different hypothalamic nuclei during development

3D analysis of OXT and AVP expression in different hypothalamic nuclei revealed a high degree of variability and autonomous specification patterns. Thus most of the rostral nuclei, particularly ADPN, BNST, PeVN and SCH, show higher degree of homogeneity (**Fig. 6a-h**) whereas caudal nuclei (SON, PVN, RCH, and AN) are more heterogeneous containing a large number of OXT+AVP neurons during early development (**Fig. 6i-p**). Among the homogeneous areas, adult ADPN and BNST are mainly formed by OXT neurons (**Fig. 6a’-b’** and **e-f**) whereas SCH is almost exclusively made up of AVP neurons^39^ (**Fig. 6d’** and **h**). Lastly, the PeVN exhibits an intermediate behavior with a low percentage of OXT+AVP neurons (**Fig. 6c’-g**). In the other hand, the heterogeneous nuclei like the PVN can be subdivided in well-defined subregions where AVP neurons concentrate at the dorsolateral (rostral view, arrow in **Fig. 6i’**) and intermediate region (lateral view, arrow in **Fig. 6m**), similarly to what has been described in the rat brain^43^. Our results revealed that this distribution is maintained from newborn to adulthood. However, at early embryonic stages OXT+AVP neurons appear mostly distributed at the rostral PVN while AVP neurons concentrate at the caudal area at E16.5 (data not shown). Similarly, OXT and AVP expression in the SON is restricted to defined areas, with a high percentage of OXT neurons at the rostral section and most AVP cells located at the ventrolateral region^43^ (**Fig. 6j’**and **n**). The RCH is also constituted by two well-differentiated areas of high cellular heterogeneity (**Fig. 6k’** and **o**), one along the surface (arrow in **Fig. 6k’**) and another surrounding the dorsomedial area (arrowhead in **Fig. 6k’**). To note, these two RCH subregions appear highly interconnected (**Supplementary video 6**), a feature that had not been appreciated previously, and that it seems to appear at early developmental stages. Among all nuclei, the AN exhibit the most pronounced cellular heterogeneity across all developmental stages since its rostral part is almost exclusively constituted by OXT and OXT+AVP neurons whereas the caudal region is formed mainly by AVP neurons (**Fig. 6l’** and **p**).

**Fig. 6.**
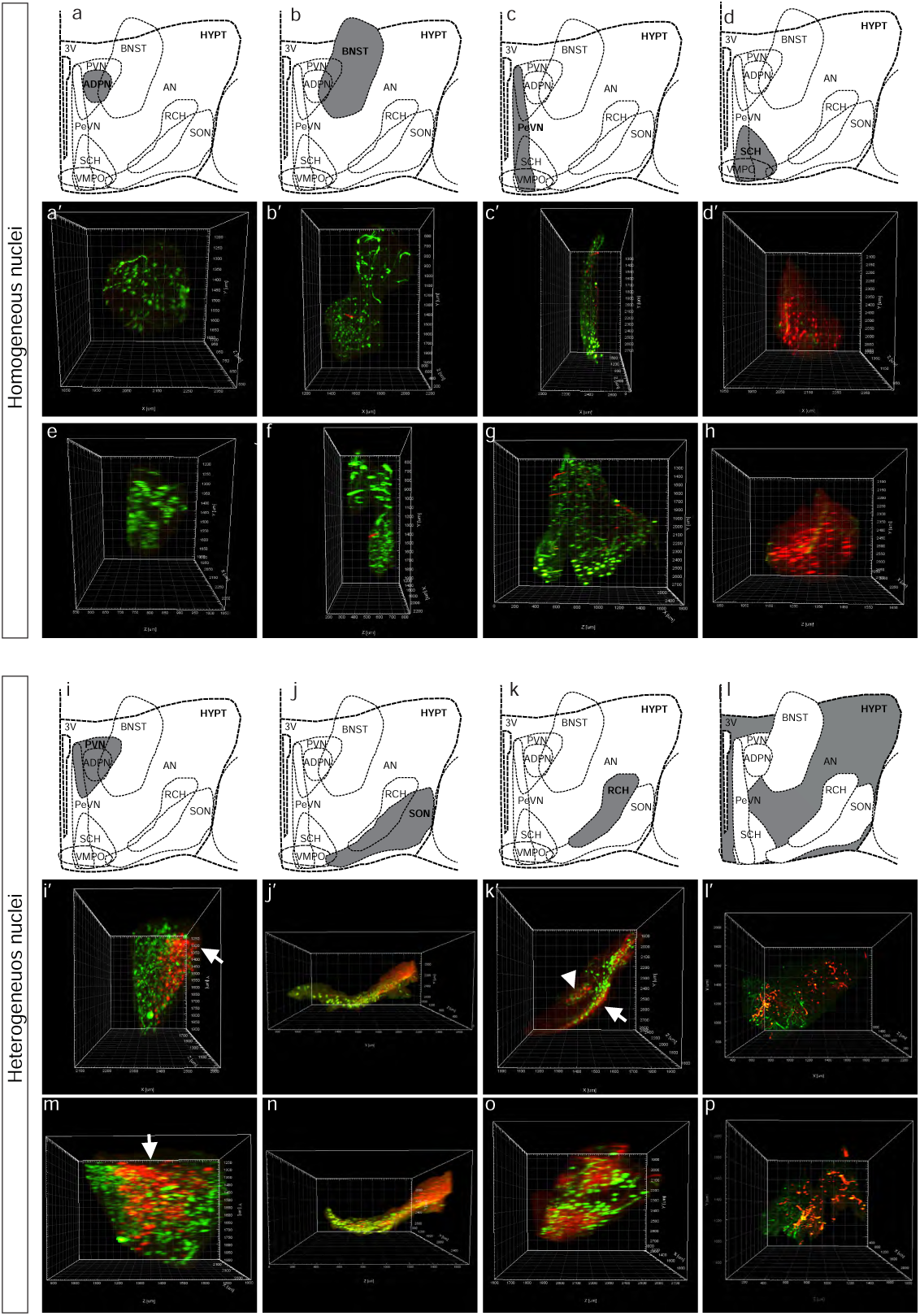
Cellular diversity within the hypothalamic nuclei. Whole-mount immunolabeling against OXT and AVP in the adult mouse hypothalamus and surrounding areas like the BNST. Nucleus segmentation shows OXT (green) and AVP (red) in different nuclei. The top panel shows the most homogeneous nuclei including ADPN, BNST, PeVN and SCH (**a-h**) and the bottom panel the most heterogeneous ones, PVN, SON, RCH and AN (**i-p**). Each nucleus (**a-d** and **i-l**) is represented by two images from different perspectives: a rostral (**a’-d’** and **i’-l’**) and a lateral view (**e-f** and **m-p**).

### Spatial distribution of OXT and AVP neurons in the PVN during development

Spatial differences in the expression pattern of OXT and AVP neurons are likely to respond to functional demands over development. We aimed to explore specific cell-type distribution dynamics analyzing the 3D reconstructions of the PVN, a prominent brain source for OXT and AVP. Despite some subtle differences, OXT and AVP neurons follow a similar expression pattern when considering their distribution from the midline and the rostro-caudal axis (**Fig. 7**). As such, during early developmental stages (E16.5, PN0), both OXT and AVP neurons can be found mostly distributed away from the midline to progressively populate the most medial regions as development progresses (**Fig. 7a-h**) but with some noticeable differences: whereas OXT neurons are more abundant in the region nearest to the midline, AVP neurons populate the intermediate regions (also see **Fig. 6m** and **6i’**). This trend is also observed in the OXT+AVP neuron population (**Fig 7. 7i-l**), which from a fairly homogeneous distribution colonize the proximal regions to the midline by PN7. In contrast, OXT and AVP expression exhibits different dynamics in the rostro-caudal axis. Young OXT and OXT+AVP neurons are abundant at the PVN caudal region to ultimately adopt an uniform expression pattern in the adult brain (**Fig 7a-d** and **7i-l**). AVP neurons in the other hand are less abundant in the caudal axis and mostly concentrate in the intermediate regions (**Fig. 7e-h**). This diversity is likely to respond to differential roles of the hypothalamic nuclei as main sources of OXT and AVP innervation, both systemically and to the CNS.

**Fig. 7.**
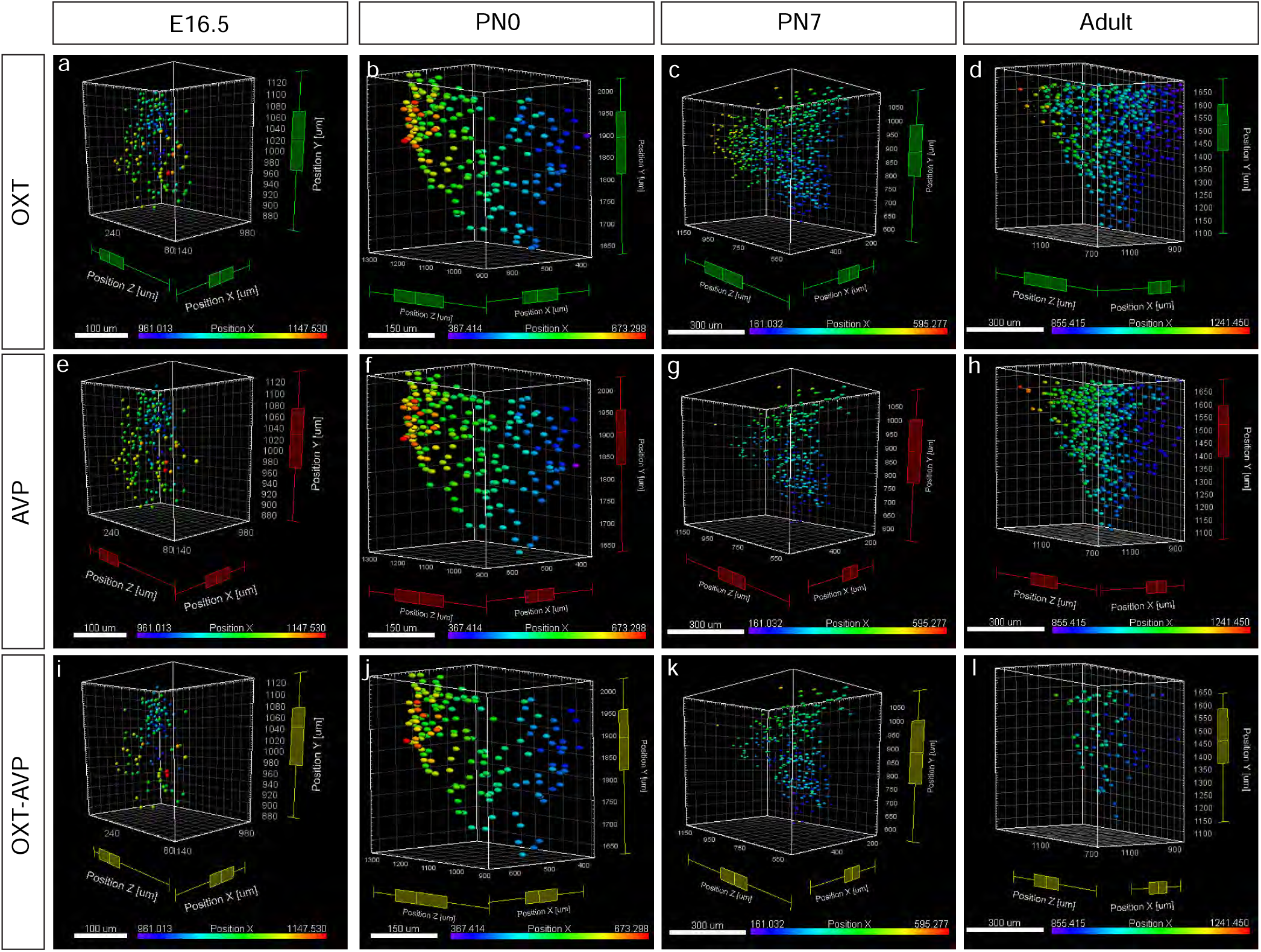
Spatial cell distribution in the PVN during development. Cell distribution of OXT (**a-d**), AVP (**e-h**) and OXT+AVP (**i-l**) neurons are represented at E16.5, PN0, PN7 and adult brain. Each plot shows the cell distribution, represented as spots in the 3D brain reconstruction. The spots color show the distance to the midline. Each developmental stage is indicated with a histogram of different color.Each plot includes the min value, quartile 1 (Q1), median, quartile 3 (Q3) and the maximum value per each axis (*x, y* and *z*). *X position*: E16.5 OXT: min = 961.00; Q1 = 1009.00; median = 1046.00; Q3 = 1072.00; max = 1146.00; E16.5 AVP: min = 972.00; Q1 = 1014.00; median = 1050.00; Q3 = 1078.00; max = 1146.00; E16.5 OXT+AVP: min = 973.00; Q1 = 1006.00; median = 1040.00; Q3 = 1069.00; max = 1147.00; P0 OXT: min = 378.00; Q1 = 467.00; median = 526.00; Q3 = 569.00; max = 673.00; P0 AVP: min = 367.00; Q1 = 477.00; median = 525.00; Q3 = 561.00; max = 673.00; P0 OXT+AVP: min = 400.00; Q1 = 477.00; median = 529.00; Q3 = 568.00; max = 673.00; P7 OXT: min = 165.00; Q1 = 258.00; median = 313.00; Q3 = 371.00; max = 536.00; P7 AVP: min = 161.00; Q1 = 257.00; median = 309.00; Q3 = 357.00; max = 595.00; P7 OXT+AVP: min = 164.00; Q1 = 263.00; median = 302.00; Q3 = 346.00; max = 514.00; Adult OXT: min = 855.00; Q1 = 915.00; median = 958.00; Q3 = 1014.00; max = 1241.00; Adult AVP: min = 859.00; Q1 = 940.00; median = 981.00; Q3 = 1026.00; max = 1230.00; Adult OXT+AVP: min = 871.00; Q1 = 934.00; median = 971.00; Q3 = 1013.00; max = 1090.00. *Z position*: E16.5 OXT: min = 89.10; Q1 = 222.00; median = 270.00; Q3 = 307.00; max = 341.00; E16.5 AVP: min = 74.00; Q1 = 236.00; median = 273.00; Q3 = 304.00; max = 341.00; E16.5 OXT+AVP: min = 88.80; Q1 = 233.00; median = 268.00; Q3 = 308.00; max = 328.00; P0 OXT: min = 890.00; Q1 = 1043.00; median = 1143.00; Q3 = 1219.00; max = 1306.00; P0 AVP: min = 891.00; Q1 = 1042.00; median = 1128.00; Q3 = 1215.00; max = 1307.00; P0 OXT+AVP: min = 892.00; Q1 = 1039.00; median = 1147.00; Q3 = 1222.00; max = 1299.00; P7 OXT: min = 504.00; Q1 = 743.00; median = 871.00; Q3 = 1002.00; max = 1162.00; P7 AVP: min = 504.00; Q1 = 747.00; median = 825.00; Q3 = 929.00; max = 1153.00; P7 OXT+AVP: min = 501.00; Q1 = 723.00; median = 802.00; Q3 = 892.00; max = 1151.00; Adult OXT: min = 669.00; Q1 = 914.00; median = 1059.00; Q3 = 1245.00; max = 1471.00; Adult AVP: min = 678.00; Q1 = 991.00; median = 1091.00; Q3 = 1198.00; max = 1468.00; Adult OXT+AVP: min = 680.00; Q1 = 978.00; median = 1069.00; Q3 = 1168.00; max = 1329.00.

### Heterogeneity of oxytocinergic cells during early development

In addition to wild type animals, we also analyzed the expression pattern of OXT in a commercial transgenic line. The OXT-Cre mice (Jackson Laboratories, ID 02423435) express Cre recombinase under the control of the OXT promoter and they are an effective tool for targeting OXT neurons^33^. To this aim, we bred these animals with a commercial tdTomato reporter line. An anti-RFP antibody was used to enhance the signal of endogenous OXT-tdTomato cells in combination with the anti-OXT antibody used in iDISCO^+^ experiments (**Fig. 8**). Analysis of the OXT-tdTomato mouse line over development revealed that OXT expression can be identified as early as E14.5 (**Fig. 8a-a’’**) with no detectable signal prior that stage (no OXT signal was detected at E12.5, data not shown). In agreement with our previous results, OXT-tdTomato cells first appear in caudal nuclei like the PVN, SON and RCH (**Figs. 8, 2** and **3**). Surprisingly, tdTomato-expressing cells do not perfectly colocalize with OXT positive neurons identified by immunohistofluorescence during embryonic and early postnatal stages. Thus, at E16.5 (**Fig. 8b-b”)** ∼ 20 % of OXT positive neurons are also RFP positive (22.81 % ± 4.44; n = 3) suggesting a high degree of heterogeneity during early development. This heterogeneity is developmentally regulated as co-labeling increases as maturation progresses (**Fig. 8c-f**) to reach an almost complete co-localization in the adult PVN (**Fig. 8f-f’’**) as previously described in this mouse line^33^ (∼ 92% co-expression in PVN and SON; https://www.jax.org/strain/024234).

**Fig. 8.**
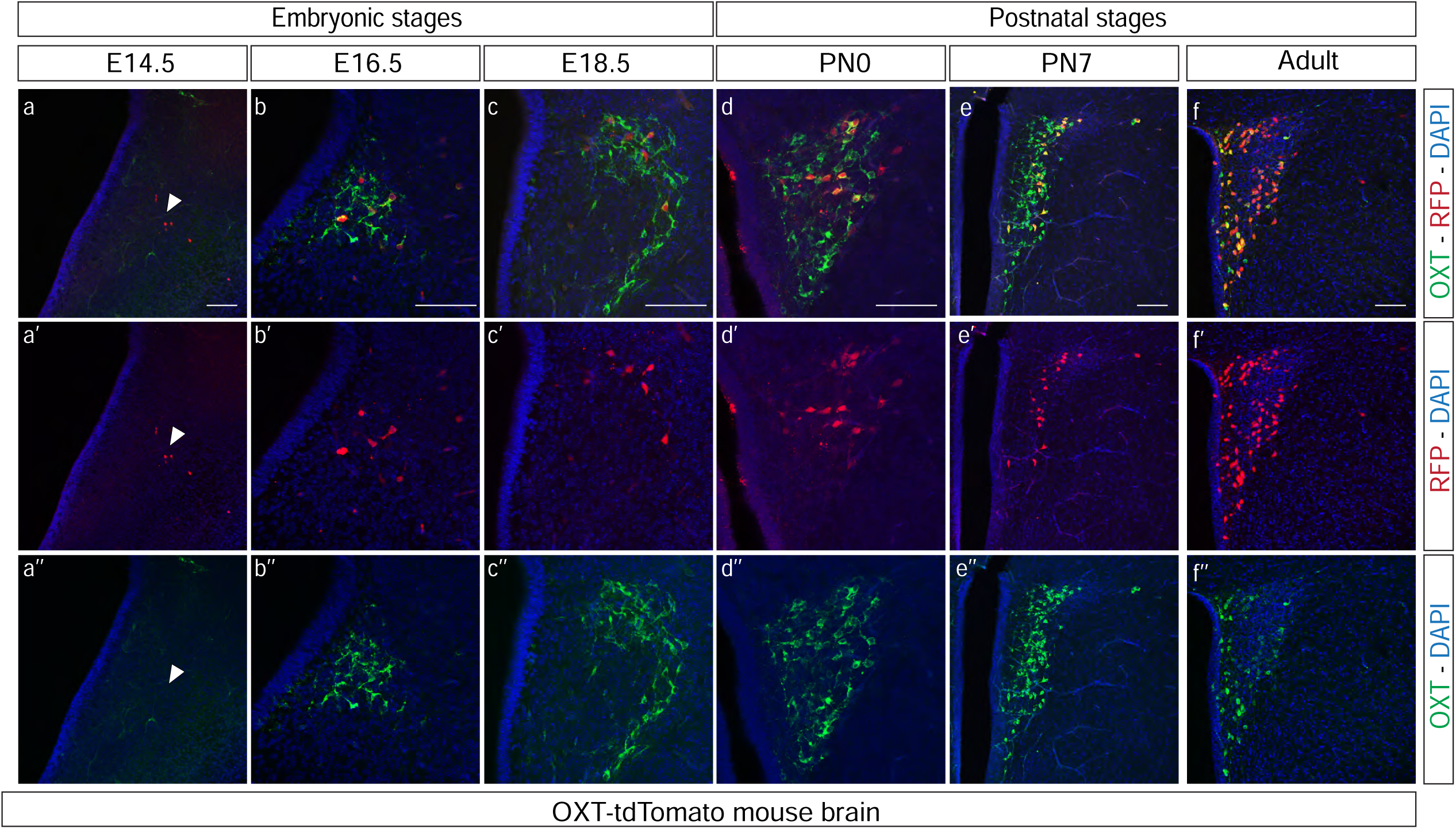
Analysis of OXT expression in an OXT-tdTomato mouse line. Coronal sections of the PVN at different stages: E14.5 (**a-a’’**), E16.5 (**b-b’’**), E18.5 (**c-c’’**), PN0 (**d-d’’**), PN7 (**e-e’’**) and adult (**f-f’’**). Immunohistofluorescence against anti-RFP (red) and anti-OXT (green). Scale bar: 50 µm in **a-d**, 150 µm in **e-f**.

The identification of a subpopulation of OXT neurons (not recognized by a common anti-OXT antibody) suggests that young oxytocinergic neurons express additional immature variants^44^ which are distinct from the ones originally characterized^45^. These differences are reduced in the adult brain, where RFP expression was seen in almost all OXT positive neurons across different nuclei. However, these early OXT variants persist until adulthood in particular nuclei like the SON and the RCH (**Fig. 9**). In fact, these nuclei retain a significant number of this oxytocinergic subpopulation at both PN7 and the adult brain, following a heterogeneous distribution in the RCH (**Fig. 9a-a’** and **b-b**) and a highly organized pattern in the SON where these neurons form a distinctive lateral subregion (**Fig. 9c-c’’** and **d-d’**). This spatial distribution in both RCH and SON reveals a niche of molecularly distinct oxytocinergic neurons that may retain some immature features in the adult brain. Our analysis reinforces the suitability of a commercial OXT-Cre mouse line to perform a detailed neuronal circuit analysis, as well as a genetic tool to study a specific subpopulation of OXT neurons in the mouse brain.

**Fig. 9.**
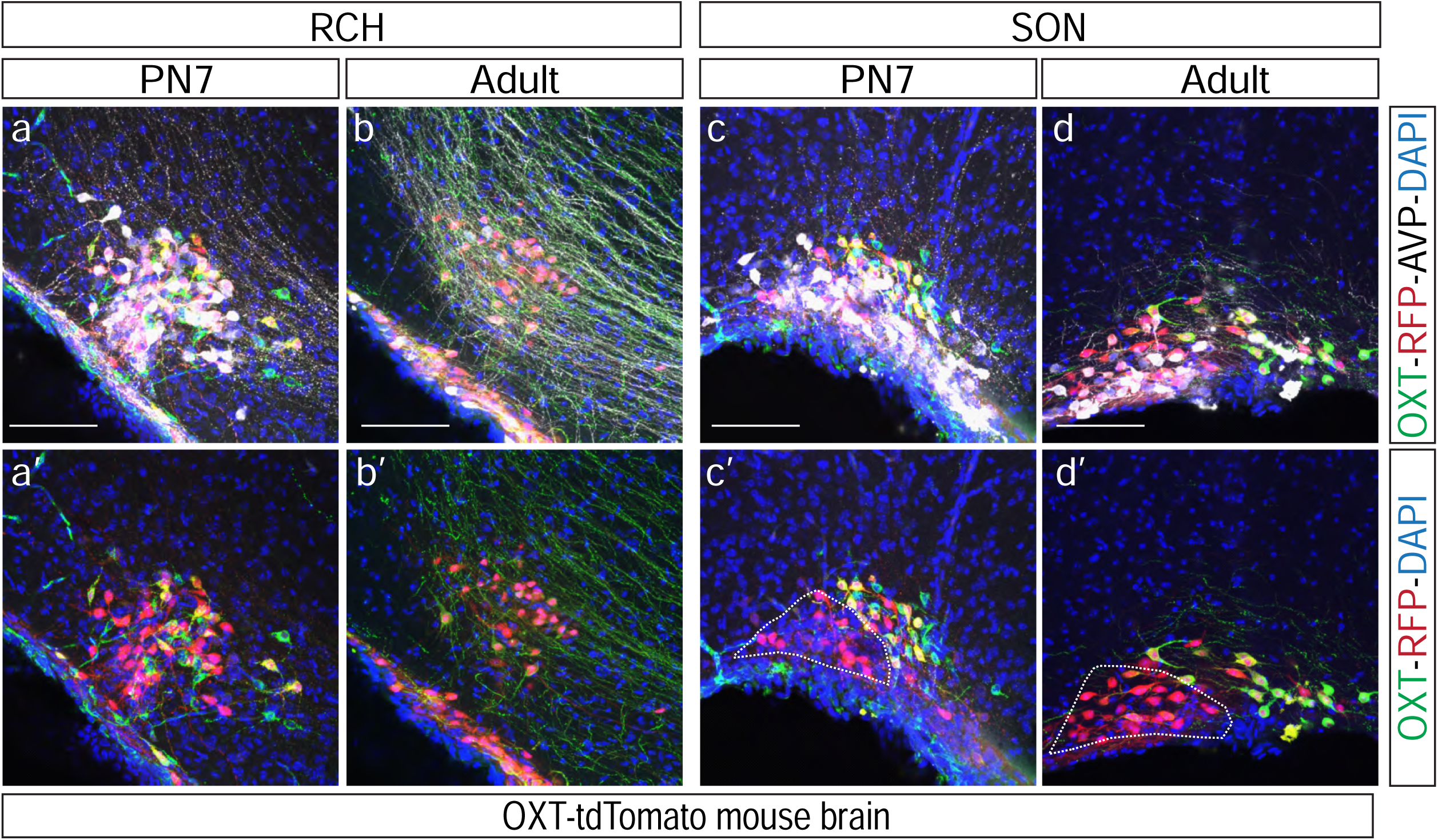
Development of SON and RCH in an OXT-tdTomato mouse line. Brain coronal sections show the RCH (**a, a’, b, b’**) and SON (**c, c’, d, d’**) at different stages: PN7 (**a, a’, c, c’**) and adult (**b, b’, d, d’**). Immunohistofluorescence against anti-OXT (green), anti-RFP (red) and anti-AVP (white). Abbreviations: SON, supraoptic nucleus; RCH, retrochiasmatic nucleus. Scale bar: 100 µm.

## Discussion

Thus far, a complete picture of the OXT and AVP innervation in the developing mouse brain was lacking in contrast to the detailed information available for the rat brain. This lack of knowledge has prevented establishing meaningful correlations between anatomical and functional data, often obtained from transgenic mouse lines and mouse behavioral studies^6,11,12,13,14,15^. Our work aims to overcome these current challenges by providing a detailed analysis of the OXT and AVP systems in the developing mouse brain. Here, we have implemented novel iDISCO^+^ clearing techniques to generate high cell-resolution brain-wide maps of the emerging and consolidated OXT and AVP neurons. Systematic 3D cell quantification revealed that the specification of these systems in the exhibit specific features distinct from the rat and other rodents which need to be considered when working with this animal model.

Similarly to other mammals^44^, mouse OXT and AVP are mostly synthesized in the PVN, SON and AN. However, vasopressinergic and oxytocinergic cells can be found in other hypothalamic areas as well as non-hypothalamic regions, where they have been much less studied. The cellular resolution achieved by the iDISCO^+^ clearing method exposed OXT and AVP neurons in non-hypothalamic areas like the MeA and, particularly the BNST (**Supplementary videos 1-5**) where OXT- and AVP-expressing neurons have been difficult to identify by traditional approaches^21-25^. These findings are in stark contrast to the rat brain where OXT and AVP neurons have not been described in the BNST regardless of the developmental stage^26-32^. These differences may relate to distinctive functions of the OXT and AVP systems in the rat and mouse brain. As such, studies in OXT knock out mice indicated that contrary to rats, OXT ablation does not prevent sexual or maternal behavior in mice although being indispensable for the milk ejection reflex and lactation^4,46^. Interestingly, the mouse BNST has been suggested to participate in social recognition through an OXT-mediated signaling involving the MeA^47^, thus the presence of a small population of OXT and AVP neurons in these areas in the mouse brain is suggestive of a local peptidergic regulation which may be unique to mice to further regulate the neuromodulatory action of long-range hypothalamic inputs.

As reported in rats, the expression of AVP and OXT can be observed at early developmental stages in the mouse PVN, SON and AN^26-32^. Interestingly, along with the rostro caudal axis, AVP neurons have been reported to appear in the dorso and ventromedial sections of the rat SON^43^ however our data indicate that the mouse brain also exhibits a well-defined pool of AVP neurons at the dorsolateral SON (**Figs. 2, 4** and **Supplementary video 2**). Moreover, the expression of OXT and AVP neurons within different hypothalamic areas revealed distinct developmental dynamics. One of the most prominent changes was observed in the SCH nucleus where the total amount of AVP neurons declined significantly over time. Previous studies have reported an increase of hypothalamic volume in parallel with organism growth^48^, thus the decrease of the relative abundance of AVP neurons in the SCH is likely to reflect a switch in the internal program of these neurons rather than overall neuronal loss due to developmentally-regulated cellular apoptosis or pruning mechanisms. SCH is considered the master regulator of the circadian clock in mammals^49^ which experiences drastic developmental changes as it transitions from non-photic cues, such as maternal hormonal cycles, to photic ones after birth^50,51^, thus we hypothesize that the developmental dynamics of SCH AVP-expressing neurons is likely to reflect this functional adaption.

Parallel to the observed changes in SCH, our findings point to a substantial reorganization of the OXT/AVP balance in most hypothalamic nuclei during development (**Figs. 2-4, Table 1** and **2**). In general terms our data indicate that the vasopressinergic system mature earlier than oxytocinergic neurons which were found scarce at PN7 in agreement with previous reports^21-25^. Immunohistochemistry and *in situ* hybridization studies in the rat brain indicated the existence of several intermediate OXT forms during early development^45^. In this regard, the analysis of a transgenic OXT mouse line (OXT-tdTomato) has revealed a significant percentage of young oxytocinergic neurons that are not recognized by a common anti-OXT antibody (**Fig. 8**) indicating the presence of early OXT variants distinct from the ones recognized by the PS38 antibody^45^ used in this study. OXT heterogeneity seems to be developmentally regulated as co-localization progressively increased over time. However, this regulation appears region-specific since areas like the SON and RCH retain a significant number of OXT-tdTomato neurons (not recognized by the PS38 anti-OXT antibody) until adulthood (**Fig. 9**). Although the existence of immature non-amidated forms of OXT in the hypothalamus is well documented^44,45^, to the best of our knowledge, this is the first report of an OXT variant specific to the SON and RCH. To note, this OXT subpopulation is organized in a particular niche within the SON suggesting the existence of a defined area with potentially distinctive properties. Our studies indicate that the OXT-Cre line employed here will be a useful tool to carry out functional analysis of this particular subpopulation of OXT neurons.

Little is known about the functional roles of immature forms of OXT and AVP peptides. OXT seems to undergo a more intensive maturation program than AVP, which has less immature variants and which mature form can be detected as early as E16.5^29^. In contrast, amidated OXT is first detected at birth coexisting with immature variants until late postnatal stages^21,32^. This phenomenon is well-documented in prairie voles where oxytocinergic cells in the PVN and SON increase from postnatal day 1–8 and 21^52^, a trend observed in the present study and suspected to be common in other mammals including humans. Intriguingly, some immature OXT forms are capable of activating the oxytocin receptor, which is widely expressed before birth^53^, suggesting a role of these variants in the consolidation of the oxytocinergic circuit. In fact, magnocellular OXT neurons exhibit undeveloped morphology and electrophysiological properties at birth which are progressively refined by reducing their spontaneous activity and decreasing action potential-evoked calcium entry during the first postnatal weeks^54,55,56^. These changes are accompanied by a switch in the regulatory mechanisms of cytosolic calcium from extrusion to sequestration into the endoplasmic reticulum^57^ which has been postulated to impact neuropeptide exocytosis^57^. In this scenario, the appearance of amidated OXT may influence the magnocellular neuron maturation as OXT modulates neuronal excitability^58^ contributing to shape synaptic transmission and plasticity. According to this notion, our analysis has revealed that the AVP and OXT circuits undergo two main maturation steps. A first one at the time of birth coinciding with a critical period for social behavior, and a second round of refinement by PN7, when the OXT/AVP ratio appears to increase in most hypothalamic nuclei. The functional consequences of this balance rearrangement is unknown, however these adjustments may underlie a critical period for hypothalamic maturation during the first two postnatal weeks^54,55,56^ which is likely involve profound cellular plasticity events as neurotransmitter switches^59^. This complex scenario is further expanded in this work by the finding of tissue-specific OXT variants (**Fig. 9**) in the SON and RCH which are likely to differentially impact the maturation of the oxytocinergic system in these particular regions and their target sites.

In summary, the present study highlights the importance of applying novel imaging tools to analyzing the properties of specific circuits in the whole brain. Improved brain clearing methods in combination with light sheet fluorescent microscopy and 3D reconstruction allows high resolution analysis of specific pathways to address their dynamics during physiological and pathological conditions. The potency of these novel technologies has revealed the developing mouse hypothalamus as a dynamic structure which undergoes intense remodeling up to the postnatal period with specific nucleus exhibiting different temporal and spatial dynamics of OXT and AVP neurons, two major candidates to regulate complex social behaviors. Our findings provide new relevant information to understand the specification of these neuronal circuits which is a critical step to uncover the wiring alterations and cellular dysfunctions underlying social deficits.

## Methods

### Animals

Experiments were performed in wild type ICR mice bred in-house. Brains were extracted at E16.5, PN0, PN7 and from young female adults (2-3 months). Oxytocin-Ires-Cre:Rosa26iDTR/+OXT-Cre (OXT-Cre thereafter) mice were obtained from Jackson Laboratories (strain ID 024234). These animals express Cre recombinase under the control of the endogenous OXT promoter and has been used previously to specifically target oxytocinergic neurons^33^. OXT-Cre mice were bred with a tdTomato reporter line (Ai9, Jackson Laboratories strain ID 007909) which exhibit a Cre reporter allele with a loxP-flanked STOP cassette preventing transcription of the red fluorescent protein variant tdTomato inserted into the Gt(ROSA)26Sor locus. OXT-tdTomato mice express robust tdTomato fluorescence following Cre-mediated recombination. All experiments were performed according to Spanish and European Union regulations regarding animal research (2010/63/EU), and the experimental procedures were approved by the Bioethical Committee at the Instituto de Neurociencias and the Consejo Superior de Investigaciones Científicas (CSIC). Animals were housed in ventilated cages in a standard pathogen-free facility, with free access to food and water on a 12 h light/dark cycle.

### Immunohistofluorescence

Mice were intracardially perfused and brains were extracted and fixed in 4 % paraformaldehyde (PFA) in PBS overnight. Embryos were fixed overnight by immersion in 4 % PFA. Samples were embedded in agarose (4 %) and sectioned at 50 µm using a Leica VT1000 S vibratome. Sections were rinsed 3 times in PBS and incubated 1 hour in PBST (0.3 % Triton, 2 % Normal Goat Serum). Samples were incubated overnight at 4 °C in a rotating shaker with the following primary antibodies: mouse anti-oxytoxin PS38^45,60^ (kindly provided by Dr. Harold Gainer; NIH); rabbit anti-RFP (Abcam, ab62341); rat anti-RFP (Chromotek, 5f8-100); rabbit anti-vasopressin (Millipore, PC234L). Tissue was rinsed 3 times in PBS and incubated with the corresponding secondary antibody: goat anti-mouse Alexa-488 (Invitrogen, A32723); donkey anti-mouse Alexa-647 (Jackson ImmunoResearch, 715-605-150); goat anti-rabbit Alexa-594 (Invitrogen, A11072); donkey anti-rat Alexa-594 (Jackson ImmunoResearch, 712-585-153). After rinsing sections were incubated 10 min at room temperature (RT) with 0.001 % DAPI (4’,6-diamidino-2-phenylindole dihydrochloride, Sigma, D9542) in PBS for nuclear staining. Slices were mounted with Mowiol 40-88 (Millipore, 475904) for histology.

### Immunolabeling-enabled three-dimensional imaging of solvent-cleared organ (iDISCO^+^)

Whole-mount immunostaining and iDISCO^+^ optical clearing procedure was performed as previously described in Renier et al., 2014 and Renier et al., 2016. Mice were intracardially perfused and brains were fixated in 4 % PFA in PBS overnight. Embryos were fixated in 4 % PFA immersion overnight. Brains were dehydrated using different concentrations of methanol (50 %, 80 %, 100 % and 100 %) until their incubation with 6 % H_2_O_2_ in methanol overnight in order to bleach the samples. After rehydratation, samples were blocked with PBS-GT (0.5 % Triton X-100, 0.2 % gelatin) during different times according to their developmental stage: overnight for E16.5 brains, 2 days for PN0 and PN7 brains, and 4 days for adult brains. Brains were incubated with OXT and AVP primary antibodies (mouse anti-oxytoxin, Gainer Lab, NIH, PS38^45,60^ and rabbit anti-vasopressin, Millipore PC234L) diluted in 0.1 % saponin (Sigma, S4521) PBS-GT solution for 5 days (E16.5), 10 days (PN0), 2 weeks (PN7) or 3 weeks (adult brain) at 37 °C on a rotating shaker. Afterwards, samples were incubated with species-specific secondary antibodies directly conjugated to a fluorophore (donkey anti-Mouse Alexa-647 (Jackson ImmunoResearch, 715-605-150); goat anti-rabbit Alexa-594 (Invitrogen, A11072) at 37 °C overnight or two days for adult brains. Immunostaining controls using secondary antibodies (without primary antibodies) were performed to test the specificity of the primary antibodies for OXT and AVP.

For the clearing, brains were dehydrated in 20 %, 40 %, 60 %, 80 % methanol in H_2_O at RT in a rotating shaker for 1 hour each step for embryo and postnatal brains, and 1.5 hours for adult brains. Then there were incubated twice in 100 % methanol for 1-1.5 hours. Samples were then treated overnight in 1 volume of 100 % methanol and l/3 volumes of 100 % dichloromethane (DCM; Sigma-Aldrich; 270997). On the next day, brains were incubated in 100 % DCM during 30 min. Lastly, samples were cleared in 100 % dibenzylether (DBE; Sigma-Aldrich; 108014) until becoming translucent.

### Microscopy and imaging processing

Confocal microscopy was performed with a laser scanning Leica SPE-II DM5550 microscope with 10×, 20× and 40× objectives. Brains were imaged on a bidirectional triple light-sheet microscope (Ultramicroscope II, LaVision Biotec) equipped with a binocular stereomicroscope (MXV10, Olympus) with a 2× objective (MVPLAPO, Olympus) using a 5.7 mm working distance dipping cap. Samples were placed in an imaging reservoir made of 100 % quartz (LaVision BioTec) filled with ethyl cinnamate (Aldrich, 112372-100G). The step size in Z-orientation between each image was fixed at 3 μm resolution for 2.5× and 4× magnifications, for nuclei identification and cell counting; and 5 μm resolution for 0.63×, 1× and 1.6× magnifications for whole brain images analysis. 3D imaging was performed using ImspectorPro software (LaVision BioTec) and processed by the Imaris x64 software (version 9.2.1, Bitplane). Video analysis was optimized for cell identification thus, providing less accurate positioning for fibers and axon terminals.

### Nuclei identification, cell counting and positioning

Quantifications of OXT and AVP cells in each hypothalamic nucleus were calculated using alternate hemispheres of four brains. Brains at different developmental stage were processed and quantified using Imaris x64 software (version 9.2.1, Bitplane) using the tools for 3D reconstruction and segmentation. Boundaries of different hypothalamic nuclei were identified according to the mouse brain atlas of Franklin and Paxinos^61^. The volume estimation of each nucleus was generated using immunohistochemical labeling as a guide and the Surface Imaris tool. False colors were given to these brain areas with the Surface Imaris tool. High-quality 3D images were selected for cell quantification. For accurate quantification of AVP- and OXT-expressing cells the fluorescence threshold was manually set and false positives were discarded of the final analysis using Imaris Spots tool. It is to note that at embryonic stages the image resolution, is not as sharp due to a greater abundance of blood vessels and neuronal processes. 3D pictures and movies were generated with Imaris. Cell 3D positioning was achieved by the statistical data-visualization module of Imaris, vantage view. High dimensional color plots were generated to analyze the 3D distribution of each cell population in the three axes (x, y and z).

### Statistics

Data were analyzed with Imaris and exported to Excel (Microsoft, Inc.) for data handling. Sample size were estimated using the power analysis set to significance level < 0.05 and 0.80 to achieve over 80 % of power with n = 4 samples per group. For statistical analysis, cell counts were estimated to follow a negative binomial distribution and adjusted to account for false positives. Results in the bar graph plot are expressed as the mean ± S.E.M. Differences were determined using an *unpaired* t-test. P < 0.05 was considered statistically significant.

## Acknowledgements

This work was supported by grants of the Spanish Agency of Research (AEI) under the grant SAF2017-82524-R (to S.J.), the “Severo Ochoa” Program for Centres of Excellence in R&D (SEV-2013-0317 and SEV-2017-0723) and the Generalitat Valenciana through the program Prometeo/2019/014 (to S.J.). The mouse anti-OXT antibody was kindly provided by Dr. Harold Gainer, NIH. We thank Dr. Cristina Marquez (IN) for her insightful comments and Dr. Juan Antonio Moreno Bravo for his help with video editing and technical issues. We also are grateful to all Jurado Lab members for their support during the realization of this work and their assistance editing the manuscript.

## Author contribution

The project was conceptualized by S. J. Brain sample preparation, data acquisition and analysis was done by M.P.M. Result interpretation, manuscript preparation and editing was done by S.J. and M.P.M. with help of members from the Jurado Lab and the Instituto de Neurociencias (CSIC-UMH).

---

**Supplementary Fig. 1.**
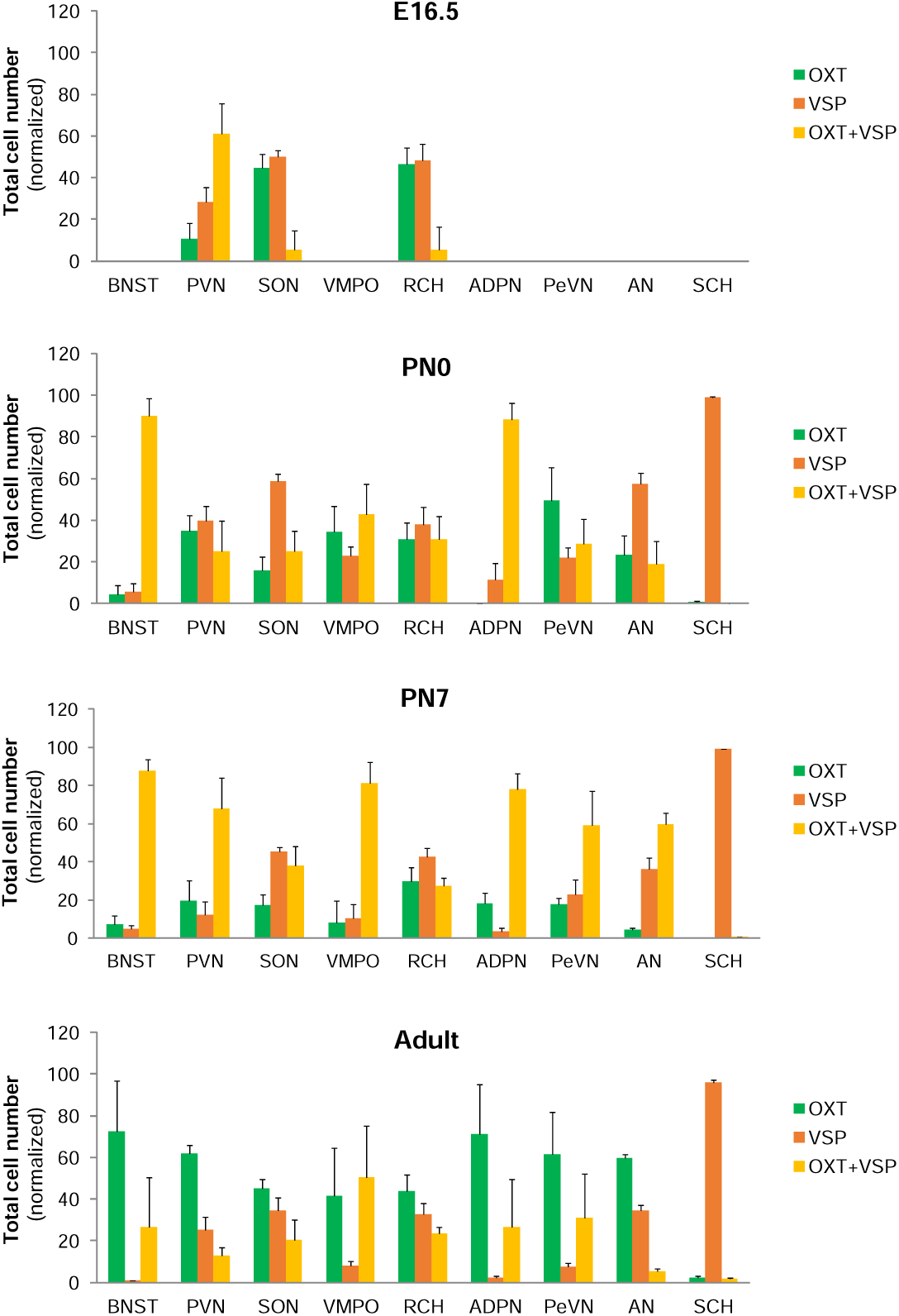
OXT and AVP dynamics during development. Percentage of OXT (green), AVP (red) and OXT+AVP neurons (yellow) are represented at E16.5, PN0, PN7 and adult brain. Data are represented as mean ± S.E.M (n = 4).

**Supplementary Fig. 2.**
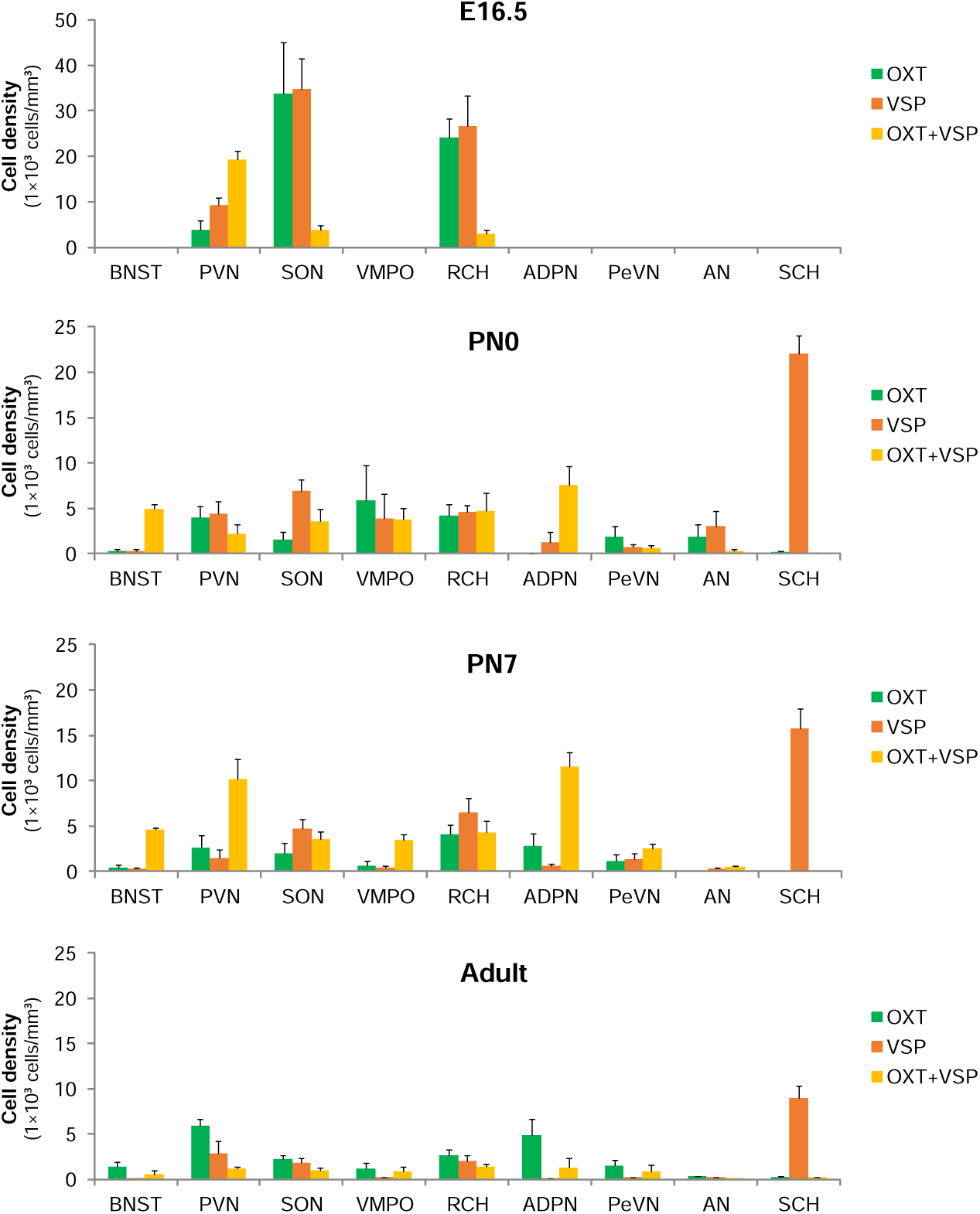
OXT and AVP expression density during development. Cell density of OXT (green), AVP (red) and OXT+AVP neurons (yellow) are shown at E16.5, PN0, PN7 and adult brain. Cell density is represented as 1×10^3^ cells/mm^3^. Data are represented as mean ± S.E.M (n = 4).

**Supplementary video 1. Identification of distinct OXT and AVP nuclei with iDISCO**^**+**^. Video shows a representative example of a cleared adult mouse brain. This technique allows to identify individual nuclei which appear labeled in different colors. Abbreviations: PeVN, periventricular nucleus; ADPN, anterodorsal preoptic nucleus; BNST, bed nucleus of strial terminalis; VMPO, ventromedial preoptic nucleus; SON, supraoptic nucleus; PVN, paraventricular nucleus; RCH, retrochiasmatic nucleus; AN, accessory nucleus; SCH, suprachiasmatic nucleus.

**Supplementary videos 2-5. 3D reconstruction of OXT and AVP nuclei in the mouse brain over development**. Whole brain immunohistofluorescence against OXT (green) and AVP (red) at different developmental stages: E16.5 (2), PN0 (3), PN7 (4) and adult (5) mouse brain. Abbreviations: PeVN, periventricular nucleus; ADPN, anterodorsal preoptic nucleus; BNST, bed nucleus of strial terminalis; VMPO, ventromedial preoptic nucleus; SON, supraoptic nucleus; PVN, paraventricular nucleus; RCH, retrochiasmatic nucleus; AN, accessory nucleus; SCH, suprachiasmatic nucleus.

**Supplementary video 6. OXT neurons interconnectivity in the RCH nucleus.** Video showing OXT neurons in the RCH area of an adult OXT-tdTomato mouse. Reconstruction of several coronal sections in the z axis revealed two well-differentiated pools of OXT neurons located at the superficial and internal RCH that exhibit high levels of connectivity.

